# Bursts of reproduction can create genetic structure in frequently recombining bacterial populations

**DOI:** 10.1101/2025.11.25.690602

**Authors:** CJ Jiang, Daniel B. Weissman

## Abstract

In many bacterial species, strong genetic structure is present, where individuals are clustered into genetically distinct groupings within the species. However, high rates of homologous recombination have also been observed in many of these species, high enough that simple models of evolution predict that such genetic structure should be eliminated. One proposed resolution to this contradiction is the presence of recurrent bursts of reproduction caused by rapid adaptation or microepidemics. We investigate this hypothesis using coalescent simulations with recombination, focusing on the distribution of pairwise genetic distances. We show that bursty reproduction can indeed create genetic structure even when recombination is so frequent that all structure would be eliminated in the absence of bursts. We find that genetic structure is only possible when there is a burst of reproduction that is sufficiently large and recent in the population’s history. In addition, we find that for other statistics beyond the distribution of pairwise distances, the simplest model of bursty reproduction does not produce distributions similar to those observed in nature. Interestingly, genetic structure from bursts of reproduction can appear among pairs of samples which do not share any genetic material by clonal descent, a feature which cannot be observed in populations whose structure is just a consequence of limited recombination.

## Introduction

Many species of bacteria exhibit stable, genetically distinct groupings, such as phylogroups in *Escherichia coli* [Clermont et al., 2000] and strains in *Staphylococcus aureus* [Raghuram et al., 2024]. What produces this within-species genetic structure? One natural possibility is that it could simply be a necessary result of clonal reproduction, with the structure reflecting the deep splits in the clonal phylogeny of the population. Equivalently, in the absence of recombination, drift produces strong linkage disequilibrium (LD) among alleles, resulting in genetic clustering of individuals. However, in some bacterial species with genetic structure, studies have inferred that recombination is very frequent relative to coalescence rates, such that a typical pair of sampled genomes will not share any genes inherited along the clonal phylogeny [Smith et al., 2000, Dixit et al., 2015], including recent studies looking at whole genome sequences [Sakoparnig et al., 2021, Liu and Good, 2024], although these results have also been challenged [Steinberg and Kussell, 2025]. Such frequent recombination should limit LD and genetic clustering, with all individuals except very close relatives roughly equally related to each other [Hanage et al., 2006]. As a result, it has been proposed that the observed deep genetic structure is the result of factors limiting genetic exchange across groups. This would be akin to the situation in sexual populations, in which genetic structure can be produced by factors such as spatial separation, genetic incompatibilities, and local adaptation. Indeed, there is strong evidence suggesting that selection does play a role in maintaining distinct groups in some species [Wiedenbeck and Cohan, 2011, Polz et al., 2013, Liu and Good, 2024]. There is also extensive evidence for molecular barriers, such as sequence similarity requirements for homologous recombination, which cause the rate of recombination to decline as genetic distance increases [Majewski and Cohan, 1999, Majewski et al., 2000, Thomas and Nielsen, 2005, Didelot et al., 2012], but it is controversial whether they are likely to have strong effects at within-species distances [Fraser et al., 2007, Dixit et al., 2017].

An alternative hypothesis for the presence of genetic structure despite frequent recombination is that it may reflect recurrent bursts of reproduction [Smith et al., 1993]. Bursts of reproduction can arise from local selective sweeps or under (micro-)epidemic population structures where infectious transmission in local environments (such as hospitals) can rapidly propagate a strain. In the past, it has been found that such microepidemic population structure can explain some aspects of genetic structure observed in *Neisseria meningitidis, Streptococcus pneumoniae*, and *S. aureus* [Smith et al., 1993, Fraser et al., 2005]. But to our knowledge, this mechanism has not been re-evaluated in the light of the much larger datasets that have become available in the last two decades.

In this work, we conduct coalescent simulations to evaluate the pattern of diversity created by bursts of reproduction. For populations without bursty reproduction, we use the standard Kingman coalescent [Wakeley, 2009]. For populations with bursty reproduction, we use the Beta coalescent. The Beta coalescent [Schweinsberg, 2003] is part of a larger family of multiple merger coalescent models that allow many lineages to coalesce at once (Fig. 1). It is especially suitable for modeling and analyzing rapidly expanding or adapting populations [Brunet and Derrida, 2012]. The Beta coalescent is of particular interest as it has been shown to describe well the phylogeny of a population where the probability of an individual having *k* or more offspring follows a power law *k*^−*α*^ [Schweinsberg, 2003]. In this work, it represents the simplest model of evolution which incorporates bursts of reproduction.

**Figure 1.**
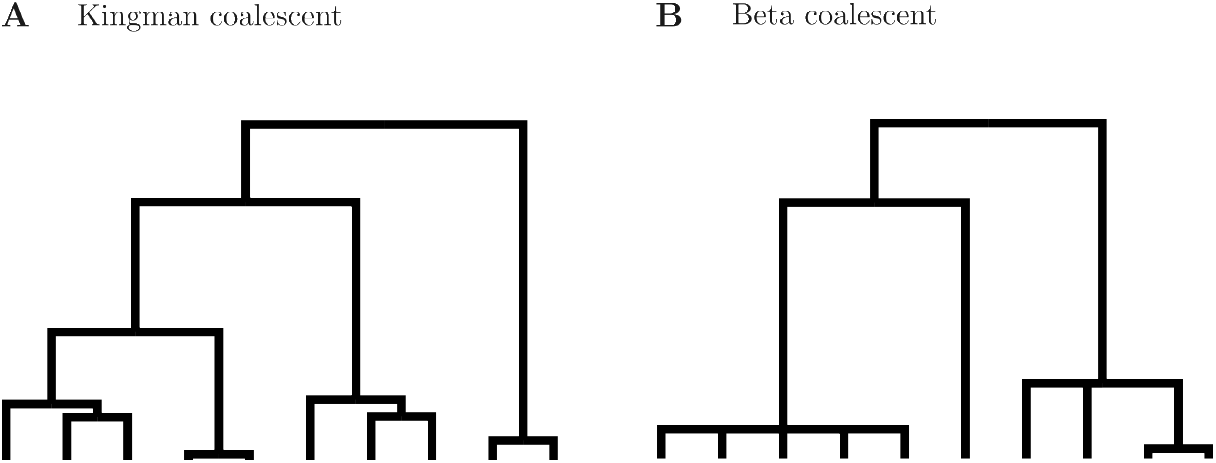
Examples of Kingman and Beta coalescent trees generated by `msprime`.

First, we demonstrate that rates of homologous recombination as high as have been inferred in natural populations are indeed sufficient to eliminate any genetic structure originating from clonal reproduction. We then show that bursts of reproduction may give rise to genetic structure even when recombination is frequent. This occurs when the burst is sufficiently large and recent in the population’s history. Even in a frequently recombining population, bursts of reproduction may lead to a temporary cluster of highly related individuals. We show that these groups can persist even after a pair of genomes has fully recombined since their most recent common ancestor. Our simulations are able to produce multi-peaked distributions of pairwise genetic distances similar to those observed in nature. However, they show distinct differences from natural populations in other diversity summary statistics, highlighting the importance of considering multiple ways of describing genetic diversity.

## Methods

### Model

We conduct coalescent simulations with recombination using msprime [Baumdicker et al., 2022]. In the Beta coalescent, the frequency and size of multiple mergers is determined by the Beta distribution *β*(2 − *α, α*) for 1 ≤ *α <* 2. When *α* → 2, the coalescent process approaches the Kingman coalescent. In our simulations, we go to the opposite extreme, setting *α* = 1.1 to allow for large multiple mergers.

Beyond the coalescent models chosen, our simulations represent the simplest possible evolutionary model. We simulate 100 samples drawn from a well-mixed population. Each sample has a linear genome of *L* = 5 Mb. We simulate homologous recombination using gene conversion [Wiuf and Hein, 2000]. Gene conversion events are initiated at rate *r* per base pair per generation and cover a geometrically distributed tract length with a mean *ℓ* = 5 kb, as observed in many gut bacteria [Liu and Good, 2024]. The key parameter governing how much clonal ancestry a typical pair of samples share is the expected number of times that a locus will have been overwritten by recombination before coalescence, *ρ* ≡ 2*E*[*T*_2_]*rℓ*, where *E*[*T*_2_] is the mean pairwise coalescence time. (*E*[*T*_2_] is often called the “effective population size”, *N*_e_, but we will use “*E*[*T*_2_]” to emphasize that for any Beta coalescent, there is no value of *N*_e_ that would produce a Kingman coalescent with the same patterns of diversity.)

Mutations are selectively neutral, meaning any rapid adaptation is occurring implicitly during a coalescent burst. Because patterns of diversity are dependent only on the ancestral recombination graph up to an overall scaling, the precise mutation rate is unimportant in our simulations. We set the mutation rate such that the expected pairwise diversity across runs is 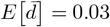, similar to natural populations, although in any given run 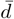 may deviate from this substantially. (We use *E*[·] to denote the expected value over different runs, and the overbar notation 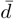 to denote the mean across all pairs within a run.) Given that we are thus fixing the mutation rate and leaving *ℓ* and, for the Beta coalecent, *α* fixed, the simulations are fully parameterized by just *ρ*.

### Pairwise genetic distances

The full genotype space is very high dimensional, necessitating the use of lower-dimensional summaries. We primarily use the distribution of pairwise genetic distances. The genetic distance *d* between a pair of samples is the fraction of bases that differ between them; 1 − *d* is the average nucleotide identity between the pair. Multiple peaks in the distribution of *d* across pairs of samples indicate the presence of genetic clusters within the population. To visually represent the full distribution of genotypes, we also project these pairwise distances onto two dimensions using metric multidimensional scaling (mMDS), specifically stress majorization using the SMACOF algorithm as implemented in <monospace>scikit-learn</monospace> [Pedregosa et al., 2011].

### Normalized index of association

We would like to have a single summary statistic which quantifies the amount of genetic structure in a population such that we can easily compare many different simulation runs. To do this, we use the normalized index of association 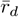, a measure of multilocus linkage disequilibrium [Agapow and Burt, 2001]. 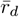 is the average correlation of pairwise genetic distances between all individuals across all loci. Specifically, if var_*j*_ is the variance of pairwise distances (between all pairs of samples) at locus *j*, and cov_*i,j*_ is the covariance of pairwise distances between loci *i* and *j*, then 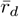 is defined as:

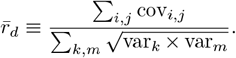

## Results

### Under the Kingman coalescent, frequent recombination always eliminates genetic structure

We first consider populations described by the Kingman coalescent. Because genetic structure arises from the structure in the clonal phylogeny, when investigating genetic structure it is natural to consider how much genetic material a pair of samples share by inheritance along that phylogeny. We classify pairs as “fully clonal” if they have experienced no recombination since their common ancestor, “partially clonal” if some regions of the genome are shared by clonal inheritance while others have recombined since the common ancestor on the clonal phylogeny, and “fully recombined” if none of the genome is shared by clonal inheritance, i.e., if every locus has been overwritten by recombination in at least one of the lineages since their common ancestor on the clonal phylogeny. We expect that fully clonal pairs will exhibit a wide range of genetic diversities *d* reflecting their idiosyncratic coalescence times. There should also be a range of *d* for the partially clonal pairs, driven by the variation in how much of their genome is clonally inherited. Fully recombined pairs, however, should have a narrow, unimodal distribution of divergence *d* peaked around the expected value *E*[*d*] = 2*E*[*T*_2_]*µ*, as the genome comprises many recombined segments with nearly independent coalescent histories [Sakoparnig et al., 2021, Liu and Good, 2024, Steinberg and Kussell, 2025]. Our simulations confirm these expectations (colored histogram bars in Fig. 2).

**Figure 2.**
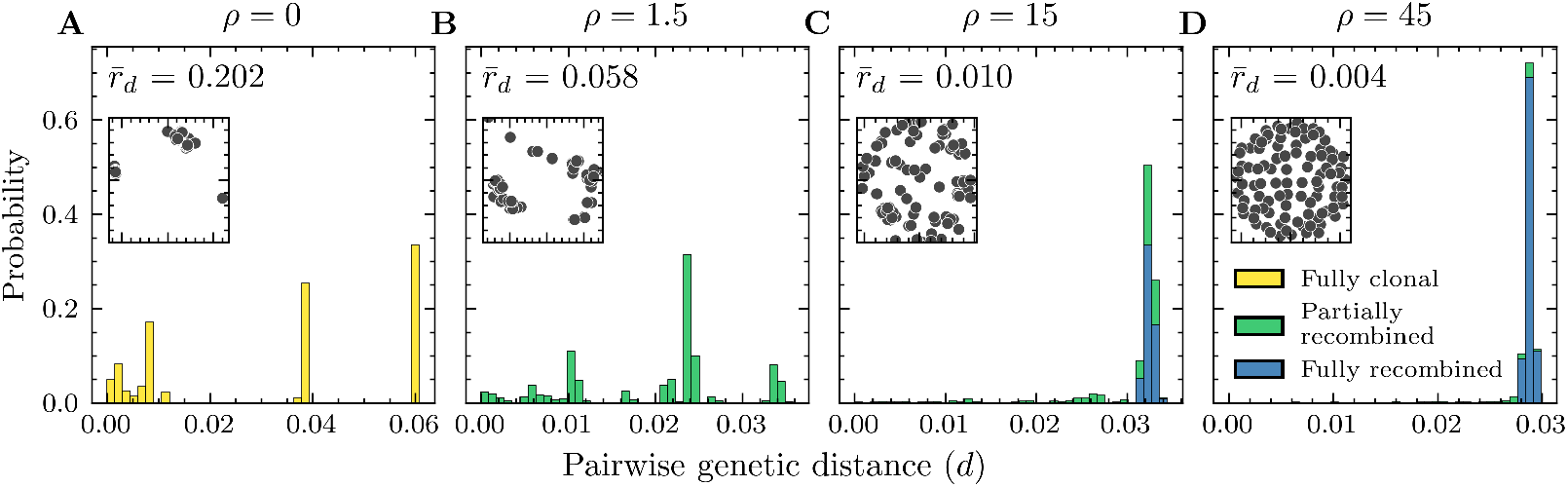
Frequent recombination eliminates genetic structure in the absence of bursty reproduction. A–D: All-vs-all pairwise distances from runs of the Kingman coalescent with varying amounts of recombination (*ρ* = 0, 1.5, 15, 45). 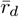 of each run is computed. Each pair in the histogram is colored based on whether all, some, or none of the genome is shared by clonal descent between the pair of samples. Insets show mMDS of samples in genotype space based on pairwise genetic distances. *ρ* = 0 and *ρ* = 1.5 show clear clustering, while *ρ* = 45 shows no clustering.

Without recombination (*ρ* = 0), genetic structure arises as a natural result of stochasticity in coalescence (Fig. 2A). The population is grouped into multiple clusters, where intra-cluster pairs are more closely related than inter-cluster pairs. At low recombination rates (*ρ* = 1.5), almost all pairs are partially recombined, and genetic structure is still present, presumably due to the clonal regions (Fig. 2B). At moderately high recombination rates (*ρ* = 15), most pairs are fully recombined or nearly so and at close to the average divergence, with only a tail of closely related partially recombined pairs creating a little structure (Fig. 2C). At very high recombination rates (*ρ* = 45), almost all pairs are fully recombined and at the average divergence (Fig. 2D). These simulations also confirm that 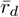 corresponds well to the notion of genetic structure and multiple peaks in the pairwise distance histogram: the loss of structure as recombination becomes more frequent is reflected in the rapid decrease in 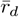 (Fig. 2).

### Bursts of reproduction can produce genetic structure even when recombination is frequent

To understand how likely genetic structure is under different recombination rates in the Kingman and Beta coalescents, we run 10^5^ replicate simulations and compute the distribution of 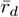. In replicate simulations without recombination, both models exhibit significant variation across runs due to the stochasticity of a single coalescent tree (Fig. 3A). We observe less LD (lower 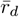) overall under the Beta coalescent than under the Kingman coalescent (Fig. 3A). This is due to an increased chance of mutations landing on external branches of the phylogeny [Eldon and Wakeley, 2008], which segregate at low frequencies, lowering LD.

**Figure 3.**
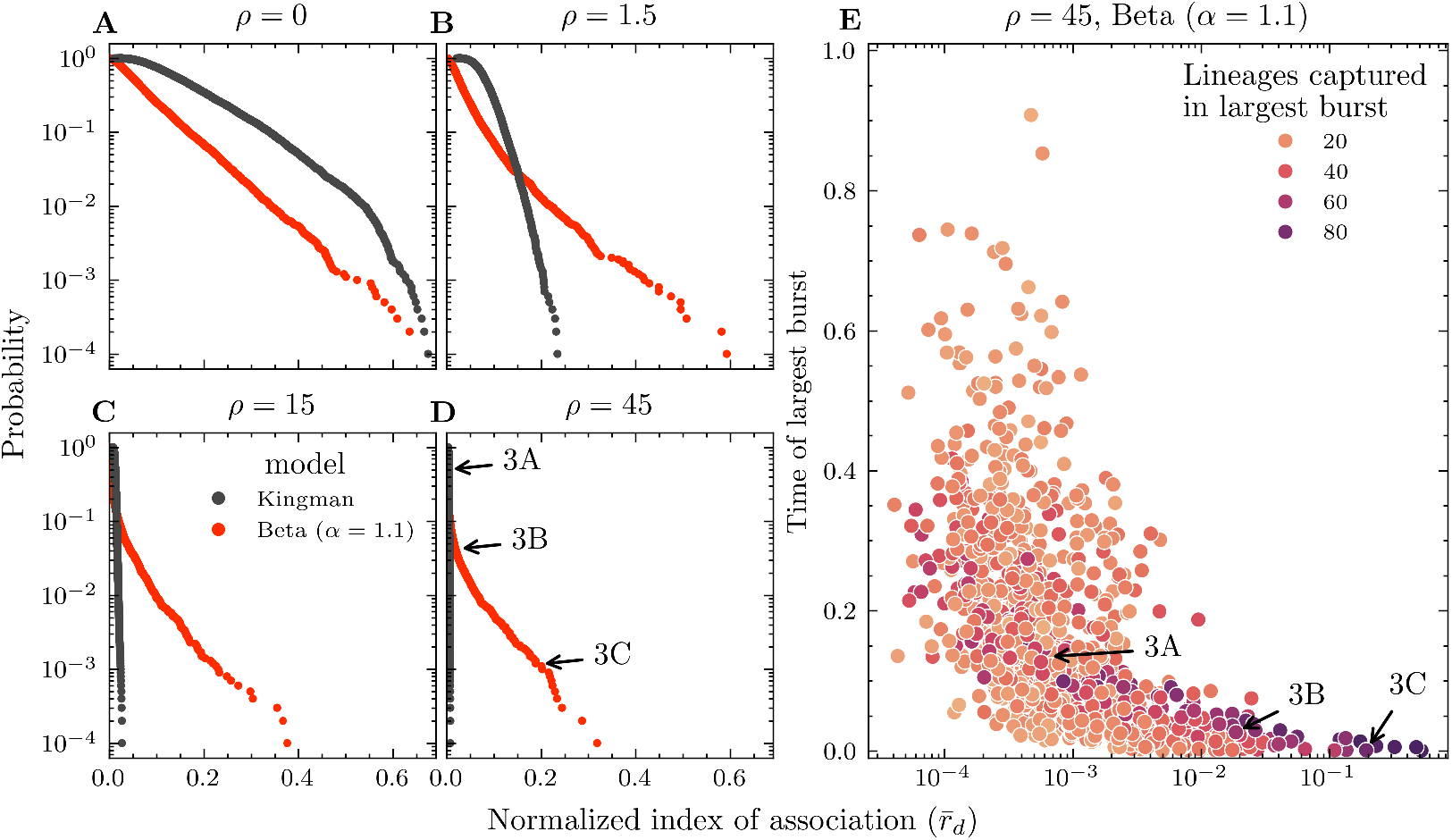
Frequent recombination completely eliminates genetic structure under the Kingman coalescent but not under the Beta coalescent. This genetic structure is only present when the burst of reproduction is both large and recent. A–D: Reverse cumulative distributions of 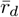 for 10^5^ replicate runs of the Kingman and Beta (*α* = 1.1) with increasing amounts of recombination (*ρ* = 0, 1.5, 15, 45). E: 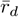 vs time *T* of the largest reproductive burst for 10^3^ replicate runs of the Beta (*α* = 1.1, *ρ* = 45). Each run is colored by the size its largest reproductive burst. Arrows in D and E point to three individual runs analyzed in Fig. 4.

When recombination is frequent (*ρ* ≥ 15), the loss of structure is essentially certain under the Kingman coalescent (Fig. 3C and D, black points), as the genome breaks up into many loci with nearly independent histories. Frequent recombination also reduces structure under the Beta coalescent, but the distribution of 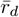 remains heavy-tailed (Fig. 3C-D, red points), meaning that some populations still exhibit structure. Specifically, runs with a large, recent burst of coalescence have high values of 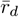 (Fig. 3E).

Interestingly, our simulations show that under the Beta coalescent, genetic structure can be present even within the subset of pairs in which no portion of the genome is inherited clonally by both partners (multiple peaks in the blue bars in Fig. 4B). This contrasts with both the intuitive argument and simulation results for the Kingman coalescent described above (Fig. 2). This feature only arises in populations that have undergone a recent large burst of coalescence. Apparently, this violates the assumption underlying the intuitive argument, that the coalescent histories of the recombined regions of the genome are nearly independent—instead, there are strong correlations in the coalescence times of the recombined regions, with some pairs having recombined regions with consistently recent coalescence times, and other pairs having recombined regions with consistently ancient coalescence times.

**Figure 4.**
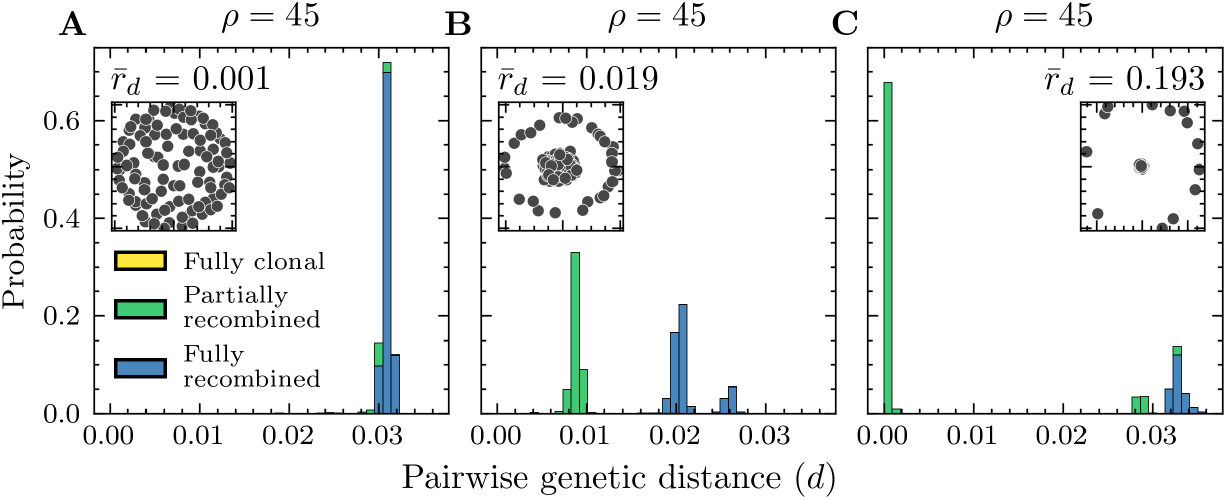
Genetic structure can appear under the Beta coalescent even when recombination is frequent. Furthermore, structure can be present within the subset of pairs which have fully recombined (B). A– C: Pairwise distance histograms of individual runs selected along the distribution of *r*_*d*_ for the Beta coalescent in Fig. 3D (*ρ* = 45). 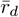 is also shown for each run. Insets show mMDS of the samples in genotype space based on pairwise genetic distances.

To understand the mechanism behind this feature, recall that a pair is considered “fully recombined” as long as they do not share any genetic material via clonal descent. Thus, at any given locus, one sample may be clonally descended from the pair’s common ancestor on the clonal phylogeny as long as the other sample has recombined at that locus. It is therefore possible for the clonal ancestry of one sample to induce correlations in the coalescence times with recombined regions of the other sample. Specifically, consider a population that experienced a large burst that coalesced a fraction *p*_coal_ of all lineages *T*_burst_ generations in the past. If the burst was sufficiently recent relative to the recombination rate, *rℓT*_burst_ ≲ 1, then at the time of the burst the ancestry of a typical sample will consist of a single lineage clonally ancestral to a large fraction *ψ* ≈ exp(−*rℓT*_burst_) = O(1) of its genome, and a large number ∼ *rLT*_burst_ of recombinant lineages ancestral to small regions of its genome. A roughly deterministic number ∼ *p*_coal_*rLT*_burst_ of the recombinant ancestral lineages will be captured by the burst of coalescence, but the single clonal lineage will stochastically be captured or escape as a whole.

This stochasticity in the fate of the clonal lineages creates three categories of pairs of samples: pairs in which both samples’ clonal ancestors were captured (Fig. 5 A and D), pairs in which one sample’s clonal ancestor was captured while the other escaped (Fig. 5 B and E), and pairs in which both samples’ clonal ancestors escaped (Fig. 5 C and F). In the first category, the pair’s clonal ancestors coalesce and the pair is therefore partially clonal. In the latter two, since at least one sample’s clonal ancestry avoids coalescence, the pair will be fully recombined. But pairs in the second category have much more coalescence occurring in the burst than those in the final category, so there will be two peaks in the divergence distribution of fully recombined pairs, with a gap between them. Table 1 lists expressions for the expected frequency of each category of pair and the fraction of the genome that typically coalesced in the recent burst for each category. We see that the genetic structure from a reproductive burst is minimal when it is too small or too ancient (and that to get all three peaks in the divergence distribution, the burst also cannot be too large or too recent).

**Table 1.** Expressions for the height and location of the three peaks in the pairwise divergence histogram produced by a large, recent burst of coalescence. *p*_coal_ is the fraction of lineages that coalesce in the burst, and *ψ* is the average fraction of the genome that is still clonally inherited at the time *T*_burst_ of the burst, *ψ* ≈ exp(−*rℓT*_burst_).

| Type of pair | Probability of pair type | Proportion of sites coalescing in burst |
| --- | --- | --- |
| Both clonal lineages coalesced at reproductive burst (Fig. 5A). | $p_{\text{coal}}^2$ | $\psi^2 + 2\psi(1 - \psi)p_{\text{coal}} + (1 - \psi)^2p_{\text{coal}}^2$ |
| Only one clonal lineage coalesced at reproductive burst (Fig. 5B). | $2p_{\text{coal}}(1 - p_{\text{coal}})$ | $\psi(1 - \psi)p_{\text{coal}} + (1 - \psi)^2p_{\text{coal}}^2$ |
| Neither clonal lineage coalesced at reproductive burst (Fig. 5C). | $(1 - p_{\text{coal}})^2$ | $(1 - \psi)^2p_{\text{coal}}^2$ |

**Figure 5.**
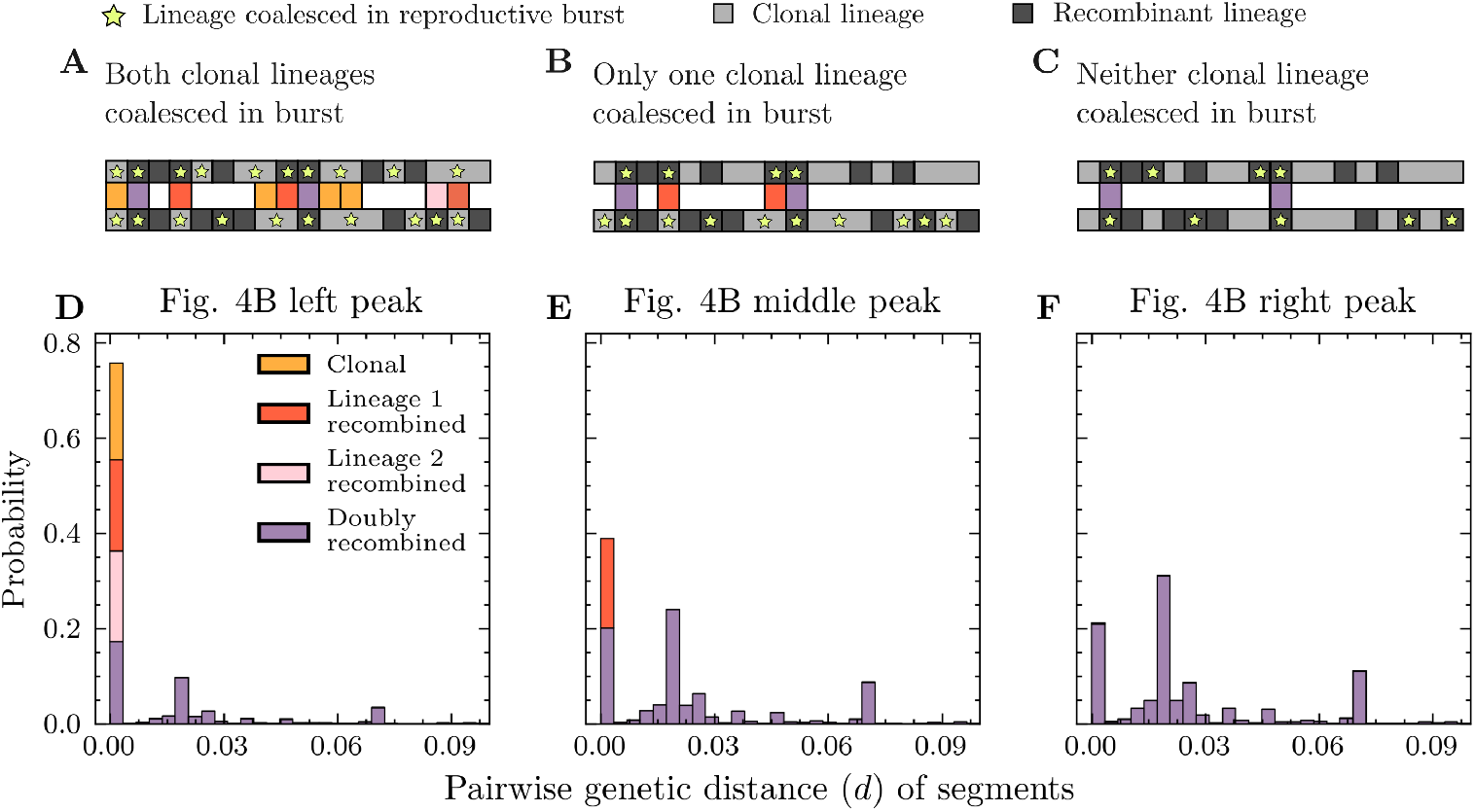
Genetic structure within fully recombined pairs occurs due to the different amounts of low divergence regions between pairs where both samples are fully recombined verses only one. A–C: Illustrations of a typical partially recombined sample, with light and dark gray rectangles representing clonal and recombinant lineages respectively. Lineages are marked with a yellow star if they coalesced at the reproductive burst. The three illustrations differ only by whether none, one, or both samples’ clonal lineages coalesced at the burst of reproduction (recombinant lineages that coalesced are the same across illustrations). At any given site, a bridge is drawn if both samples coalesced at the burst of reproduction and colored based on the relationship between the two samples at that site. D–F: Distribution of *d* for segments between all pairs for each of the three peaks in Fig. 4B. A segment is a contiguous block of sites that all share the same ancestry. Segments are colored depending on if the pair is clonal (neither lineage is recombinant), singly recombined (one lineage is recombinant), or doubly recombined (both lineages are recombinant) at that segment.

### Frequency of identical blocks of genome

One shortcoming of pairwise genetic distances is that it does not differentiate between diversity introduced by recombination verses mutation. If diversity is introduced primarily by mutations accumulating on a clonal lineage, differences should be distributed approximately uniformly across the genome. If diversity is primarily introduced by recombination, differences should instead be concentrated in short stretches of genome. The comparison between the fraction *f* of identical *l* = 1 kb blocks across the genome and genetic distance *d* can quantify the degree to which a population follows either of these regimes [Liu and Good, 2024].

First, we unify Liu and Good [2024]’s expressions for the expectation of *d* given *f* under the assumption that recombinant sequences are well-approximated as being of average divergence 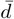. Consider a pair of samples, and suppose that the time to their most recent common ancestor in the clonal phylogeny is *T*_2_. Note that they may not have inherited any genetic material from this ancestor. Each block of length *l* in a lineage is replaced at a rate *R* ≈ *rℓ* exp(− *l/ℓ*), where *ℓ* is the typical recombination tract length. In our simulations, *ℓ* = 5 kb is large compared to *l* = 1 kb, so *R* ≈ *rℓ*—if a recombination event is initiated within *ℓ* bases of a block, it is likely to replace the block. The expected fraction of blocks that have been clonally inherited in both lineages from the most recent common clonal ancestor is therefore P(no recombination) = exp(−2*RT*_2_). Assuming recombinant segments are of average divergence 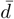, the expected divergence *d* given *T*_2_ is then:

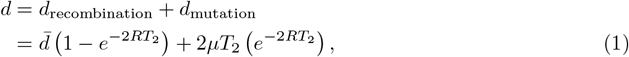

where *µ* is the per-base mutation rate. The expected fraction *f* of identical blocks given *T*_2_ is set by the probability that a block avoided both recombination and mutation:

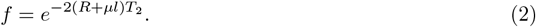

Combining (1) and (2), we can eliminate *T*_2_ and directly relate the expected divergence *d* and the expected fraction of identical blocks *f* :

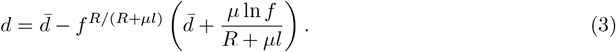

(3) accurately describes the relationship between *d* and *f* under the Kingman coalescent regardless of the recombination rate (Fig. 6A). However, for the Beta coalescent, in the moderately structured simulation run shown in Fig. 4B, the pattern of identical blocks created by reproductive bursts does not follow this expectation. Instead, even very diverged pairs, including fully recombined pairs, can share *f* ∼ 10% identical blocks, when (3) would predict that they would have almost none. (3) fails in this case because the assumption that recombined blocks have close to the average divergence breaks down: as explained above (Fig. 5F), a substantial fraction of recombined blocks will coalescence in the recent burst of coalescence, and so can be identical and contribute to *f*.

**Figure 6.**
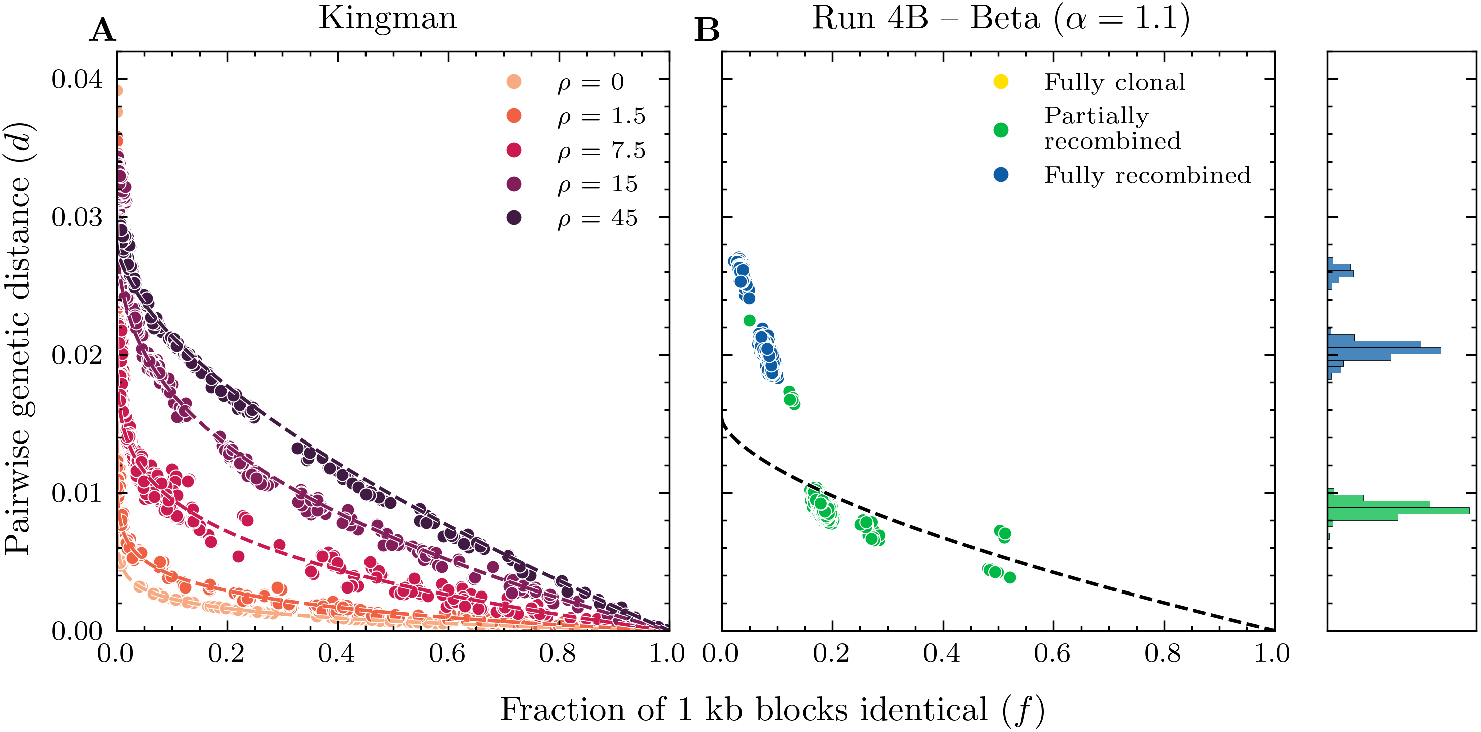
The pattern of diversity by reproductive bursts does not adhere to the expectation under an assumption of average divergence recombinant segments. Simulations of the Kingman do follow this expectation. A: Pairwise distances *d* vs. fraction of 1 kb blocks that are identical between all pairs in simulations of the Kingman under different recombination rates (*ρ* = 0, 1.5, 7.5, 15, 45). Dotted lines show expectation from model assuming average divergence recombination for each recombination rate. B: Analogous version of A for the Beta run (*α* = 1.1, *ρ* = 45) from Fig. 4B. The histogram on the right shows the same distribution of pairwise distances as in Fig. 4B.

Liu and Good [2024] use the fraction of identical blocks *f* between a pair of samples as a proxy for the fraction of the genome that is clonally inherited by both from their most recent common ancestor. We see that this is a poor approximation when bursts of reproduction occur. Even in pairs where both samples have completely recombined, two samples share many identical blocks. This is because while neither samples’ clonal lineage was part of the reproductive burst, both recombined with individuals who were, resulting in regions of the genome where they are highly related (Fig. 5F).

We note that this deviation from (3) is intrinsic to the simplest model of bursty reproduction. There are no alternative parameters (varying block size or mutation rate) or burst timing and size which eliminates this deviation whilst preserving genetic structure. Subsequently, we conclude that our model fails to match the pattern of identical blocks in Liu and Good [2024], where many species have structure even within pairs that share no identical blocks. However, we can imagine more fine-tuned models of bursty reproduction, where for example samples coalesce in smaller bursts before coalescing in the biggest burst, may better reduce the deviation from this expectation.

## Discussion

The recent influx of whole-genome sequences across numerous bacterial species has called into question simple models of bacterial evolution and recombination [Hanage, 2016]. Most notably, many species with genetic structure appear to recombine frequently enough that this structure should seemingly be erased [Fraser et al., 2007, Sakoparnig et al., 2021, Liu and Good, 2024]. There are many possible explanations for this phenomenon, including that the frequency of recombination has been overestimated [Steinberg and Kussell, 2025]. Assuming however that recombination is frequent, most explanations can be seen as invoking some form of non-uniform barriers to recombination, where some sets of individuals are more likely to exchange genetic material than others. The exception is the idea that structure has been generated simply by bursts of reproduction [Smith et al., 1993, Fraser et al., 2005]. This explanation preserves the exchangeability of lineages, and therefore is particularly tractable for model. In this work, we simulated the simplest model of bursty reproduction, the Beta coalescent, to test its ability to match patterns of genetic structure found in nature. We found that bursts of reproduction can indeed produce genetic structure in a population even when recombination is frequent. However, genetic structure is only noticeable when the reproductive burst is unusually recent and large. This simple model could thus only explain the presence of structure in a small minority of populations, not as a frequent occurrence across many populations as is observed in nature [Rodriguez-R et al., 2024, Liu and Good, 2024].

Surprisingly, we also find that bursty reproduction can produce genetic structure within the subset of pairs which have fully recombined since their most recent common ancestor in the clonal phylogeny, i.e., which share no genetic material via clonal descent. One way to understand this intuitively is to note that at any given locus, one member of the pair may still be clonally descended from the common ancestor as long as the other is not, so it is still possible for the clonal phylogeny to influence statistics of the genetic diversity. This issue may become even more relevant when considering, e.g., trees relating many samples, as in Sakoparnig et al. [2021]. As long as just one sample has clonal ancestry, it is still conceivable that the clonal phylogeny could be reflected in the statistics of the trees, although it is not clear from our results whether this effect is likely to be significant.

Here we considered only the simplest possible model of bursty reproduction, so while it is unable to match all aspects of naturally observed genetic diversity, other models of bursty reproduction may be closer. While the Beta coalescent is a popular model for simulating peculiar life histories and rapid adaptation, many others have been proposed [Lin and Kussell, 2017, Tellier and Lemaire, 2014]. Most notably, the Ξ-coalescent, which allows for simultaneous multiple mergers that could represent regular extinction and recolonization events, could provide a better match to data [Schweinsberg, 2000].

Beyond the presence of genetic structure, it is also surprising that the genetic distance boundaries which define intra-species units have been found to be similar across many different species [Rodriguez-R et al., 2024]. For many human-associated bacteria, such as pathogenic and commensal bacteria or those which inhabit the gut microbiome, these natural boundaries may reflect the corresponding rapid expansion in human populations over the last few centuries. Bacterial species which rapidly expand following the expansion of their host human populations will share bursts of reproduction at similar times, creating similar patterns of genetic structure. As such, simulations which account for a variable population size by increasing coalescence rates closer to the present may also provide a better fit.

As the evolution of bacteria in nature is likely determined by not one but many of the hypotheses proposed, it may be necessary to combine these hypotheses in future models to accurately reflect natural patterns – such as useful population structure, complex fitness landscapes, and biased recombination. However, as we depart from these simple models which represent the extremes of the evolutionary parameter space, issues regarding how to parameterize these complex models or what to use as a meaningful null become a difficult problem [Sakoparnig et al., 2021]. Future work should aim to resolve these uncertainties and provide reasonable modeling frameworks that are both tractable and accurate in describing our growing understanding of bacterial evolution.

## Acknowledgements

The authors thank Franz Baumdicker and Peter Ralph for helpful conversations.

